# Preclinical Development of a Stabilized RH5 Virus-Like Particle Vaccine that Induces Improved Anti-Malarial Antibodies

**DOI:** 10.1101/2024.01.04.574181

**Authors:** Lloyd D. W. King, David Pulido, Jordan R. Barrett, Hannah Davies, Doris Quinkert, Amelia M. Lias, Sarah E. Silk, David J. Pattinson, Ababacar Diouf, Barnabas G. Williams, Kirsty McHugh, Ana Rodrigues, Cassandra A. Rigby, Veronica Strazza, Jonathan Suurbaar, Chloe Rees-Spear, Rebecca A. Dabbs, Andrew S. Ishizuka, Yu Zhou, Gaurav Gupta, Jing Jin, Yuanyuan Li, Cecilia Carnrot, Angela M. Minassian, Ivan Campeotto, Sarel J. Fleishman, Amy R. Noe, Randall S. MacGill, C. Richter King, Ashley J. Birkett, Lorraine A. Soisson, Carole A. Long, Kazutoyo Miura, Rebecca Ashfield, Katherine Skinner, Mark Howarth, Sumi Biswas, Simon J. Draper

## Abstract

The development of a highly effective vaccine against the pathogenic blood-stage infection of human malaria will require a delivery platform that can induce an antibody response of both maximal quantity and functional quality. One strategy to achieve this includes presenting antigens to the immune system on virus-like particles (VLPs). Here we sought to improve the design and delivery of the blood-stage *Plasmodium falciparum* reticulocyte-binding protein homolog 5 (RH5) antigen, which is currently in a Phase 2 clinical trial as a full-length soluble protein-in-adjuvant vaccine candidate called RH5.1/Matrix-M™. We identify disordered regions of the full-length RH5 molecule induce non-growth inhibitory antibodies in human vaccinees, and a re-engineered and stabilized immunogen that includes just the alpha-helical core of RH5 induces a qualitatively superior growth-inhibitory antibody response in rats vaccinated with this protein formulated in Matrix-M™ adjuvant. In parallel, bioconjugation of this new immunogen, termed “RH5.2”, to hepatitis B surface antigen VLPs using the “plug-and-display” SpyTag-SpyCatcher platform technology also enabled superior quantitative antibody immunogenicity over soluble antigen/adjuvant in vaccinated mice and rats. These studies identify a new blood-stage malaria vaccine candidate that may improve upon the current leading soluble protein vaccine candidate RH5.1/Matrix-M™. The RH5.2-VLP/Matrix-M™ vaccine candidate is now under evaluation in Phase 1a/b clinical trials.

## Introduction

Over the past two decades, the number of deaths from malaria, caused by the *Plasmodium falciparum* parasite, has been steadily declining due to the improved deployment of antimalarial tools. However, the success of malaria control measures requires sustained investment, which is expensive and threatened by the emergence of drug and insecticide resistance. Moreover, worrying evidence suggests progress has stalled in recent years, with malaria cases and deaths rising since 2019 ^1^. Hence, there remains an urgent need for the development of transformative new tools, including highly efficacious and durable malaria vaccines, to complement and/or replace current malaria prevention public health measures. Substantial recent progress has been made in this area, with the RTS,S/AS01 (Mosquirix™) and R21/Matrix-M™ subunit vaccines (that both target the circumsporozoite protein [CSP] on the liver-invasive sporozoite stage of *P. falciparum*) showing efficacy against clinical malaria in young African infants ^2,3^. However, efficacy wanes over time and if a single sporozoite slips through the net of protective immunity and infects the liver, then the subsequent disease-causing blood-stage of infection is initiated. Seasonal vaccination has been demonstrated to be highly efficacious in Phase 3 trials, with RTS,S/AS01 (Mosquirix™) non-inferior to seasonal malarial chemoprophylaxis (SMC), which has been associated with approximately 75% efficacy ^4–6^; however, annual vaccination is expensive and a major burden on already stretched health systems ^7^. Vaccination against the blood-stage merozoite, aiming to prevent erythrocyte invasion and the clinical manifestation of malaria disease, represents an alternative and complementary approach. Moreover, the combination of a new blood-stage anti-merozoite vaccine with existing anti-sporozoite vaccines is currently regarded as a leading future vaccination strategy to achieve higher and more durable efficacy ^8^.

Merozoite invasion of human erythrocytes occurs rapidly and in a complex multistep process requiring numerous parasite ligand-host receptor interactions. Historic blood-stage vaccine candidates struggled because many of these parasite ligands are highly polymorphic and the interactions they mediate are redundant ^9^. However, the identification of the reticulocyte-binding protein homolog 5 (RH5) ^10^ has renewed vigor in the *P. falciparum* blood-stage vaccine field over the last decade ^8^. RH5 is an essential, highly conserved and antibody-susceptible antigen, delivered to the parasite surface in a pentameric protein complex ^11–13^ where it binds to host basigin/CD147 ^14^. This receptor-ligand interaction is critical for parasite invasion ^15^, underlies the human host tropism of *P. falciparum* ^16^, and vaccination of *Aotus* monkeys with RH5 conferred significant *in vivo* protection against a stringent blood-stage *P. falciparum* challenge ^17^. These preclinical data supported onward progression of RH5-based vaccine candidates to the clinic, with four early-phase clinical trials now completed in the UK or Tanzania; each of these studies utilized vaccines that deliver the full-length RH5 molecule (RH5_FL) using either a viral-vectored platform ^18,19^ or a recombinant protein called RH5.1 ^20^ formulated in AS01_B_ adjuvant from GSK ^21^ or Matrix-M™ adjuvant from Novavax (ClinicalTrials.gov NCT04318002). All of these vaccines have shown acceptable safety and reactogenicity profiles, with the highest levels of antibody observed when using the protein-in-adjuvant formulations ^21^ and/or when vaccinating Tanzanian infants as opposed to UK or Tanzanian adults ^19^. The RH5.1/Matrix-M™ vaccine candidate has since progressed to a Phase 2b field efficacy trial in 5-17 month old infants in Burkina Faso (ClinicalTrials.gov NCT05790889).

All of these RH5-based vaccine candidates have induced serum IgG antibodies in humans that mediate functional growth inhibition activity (GIA) against *P. falciparum in vitro*. Notably, despite differences in the quantity of anti-RH5 serum IgG induced, all of these vaccine candidates tested to-date show comparable functional quality of the anti-RH5 human IgG ^19^, i.e., they achieve the same amount of GIA *in vitro* per unit of anti-RH5 antibody, consistent with all vaccine candidates encoding almost identical immunogens based on RH5_FL. Importantly, functional antibody activity, as measured using the *in vitro* assay of GIA, has also been shown to correlate with efficacy against experimental *P. falciparum* blood-stage challenge of both *Aotus* monkeys ^17^ and UK adults ^21^. This vaccine-induced mechanism of protection against blood-stage *P. falciparum* was subsequently validated by passive transfer of anti-RH5 monoclonal antibody (mAb) in *Aotus* monkeys ^22^ and a humanized mouse model ^23^. However, despite this progress, the overall quantity of anti-RH5_FL IgG associated with protection in the *Aotus* monkey model was high ^17^. We therefore sought here to develop an improved RH5-based vaccine candidate that could substantially outperform the current clinical lead vaccine candidate, RH5.1/Matrix-M™, in terms of quantitative and/or qualitative antibody immunogenicity. To do this, we explored rational re-design of the RH5 immunogen based on serological analyses of the anti-RH5.1 IgG from clinical trials and improved delivery of RH5 using a virus-like particle (VLP) platform. In the case of the latter, given the well described challenges of recombinant RH5 protein expression, we elected to test a “plug-and-display” strategy using SpyTag-SpyCatcher bioconjugation technology ^24,25^. We also elected to use the hepatitis B surface antigen (HBsAg) VLP scaffold ^26^, given the extensive safety track record of the hepatitis B vaccine and to align the delivery platform with that used for delivery of the CSP antigen by both RTS,S and R21 ^2,3^.

## Results

### Vaccine-induced human anti-RH5 growth inhibitory antibodies target RH5ΔNLC

We previously assessed the RH5.1/AS01_B_ vaccine candidate in healthy malaria-naïve UK adults, using a variety of dosing and immunization regimens ^21^. The RH5.1 protein was manufactured in a *Drosophila* Schneider 2 (S2) stable cell line system and comprises the whole ∼60 kDa RH5 soluble molecule with four sites of potential N-linked glycosylation removed ^20^. This molecule therefore includes the structured alpha-helical core of RH5 (termed “RH5ΔNLC”) and the predicted regions of disorder: the long N-terminal region, intrinsic loop and small C-terminus (**Fig. 1A**). The structure of the α-helical core protein (including the small C-terminus but lacking the N-terminus and intrinsic loop, known as “RH5ΔNL”) was previously reported ^27^. Human serum samples, collected after three immunizations with RH5.1/AS01_B_, were all positive for IgG by ELISA against the recombinant full-length RH5.1, RH5 N-terminus (RH5-Nt) and RH5ΔNL proteins. Responses were comparable and did not differ significantly by vaccine dose or delivery regimen (**Fig. 1B**). Sera were also tested by ELISA against a linear peptide array spanning the RH5.1 antigen sequence. Responses were clearly detectable across all the regions of predicted protein disorder (N-terminal region, intrinsic loop and C-terminus), confirming these contain linear antibody epitopes which appear largely absent in the α-helical core regions (**Fig. 1C**). Vaccine-induced anti-RH5.1 serum IgG responses thus reacted across the whole molecule, including regions comprising both linear and conformational epitopes.

**Figure 1.**
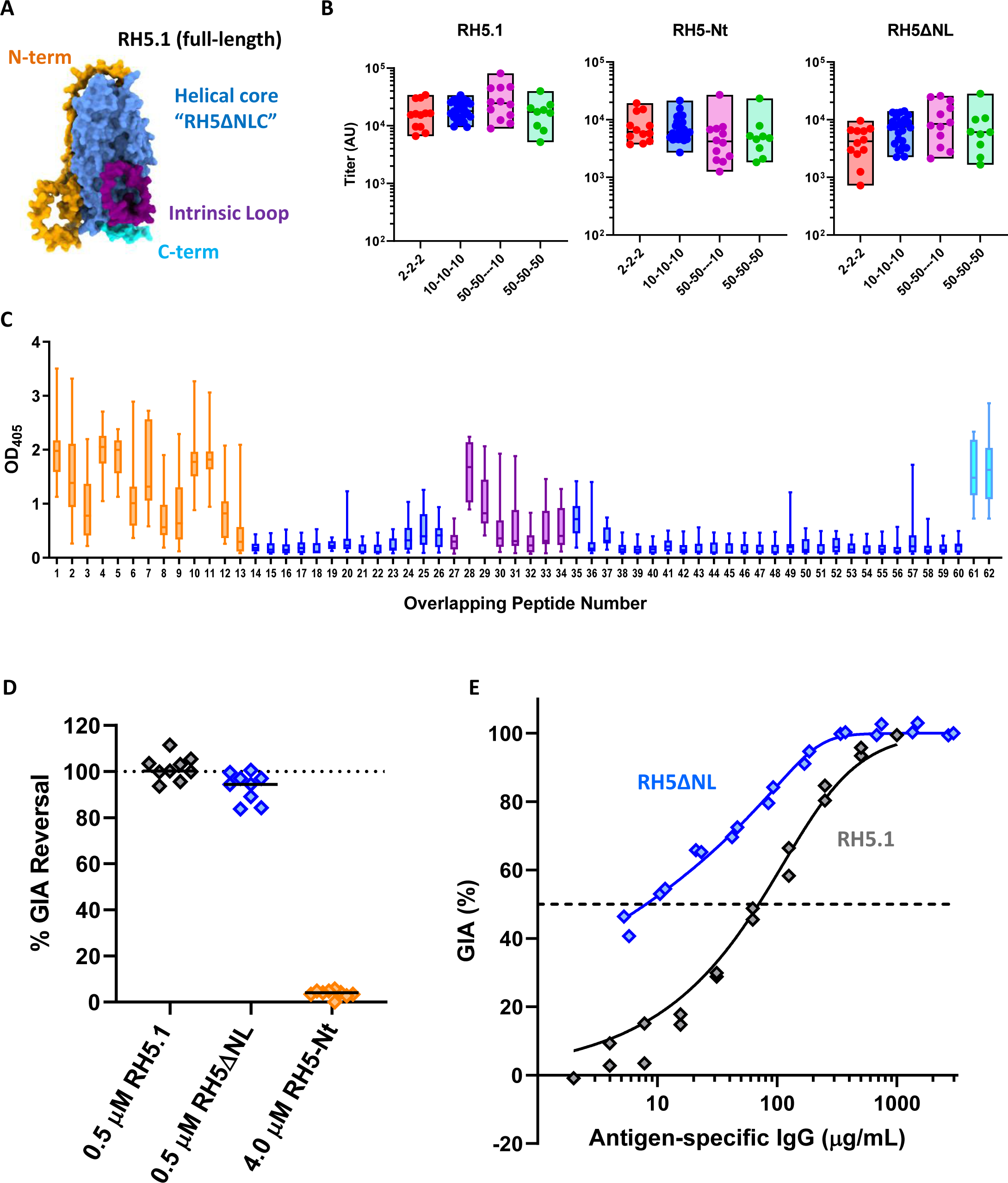
Assessment of vaccine-induced human anti-RH5.1 antibody targets. (**A**) AlphaFold model (#AF-Q8IFM5-F1) of the full-length RH5 molecule on which the RH5.1 protein (amino acids [aa] E26-Q526) vaccine ^20^ was based. The structured alpha-helical core (“RH5ΔNLC”) is shown in light blue, whilst the regions of predicted disorder include: i) the linear N-terminus (RH5-Nt; aa E26-Y139; orange); the intrinsic loop (aa N248-M296; purple); and small C-terminus (aa D507-Q526; cyan) ^27^. (**B**) Serum IgG antibody titers in RH5.1/AS01_B_ vaccinees as measured by ELISA against recombinant RH5.1, RH5-Nt and RH5ΔNL proteins in arbitrary units (AU). Vaccinees received three doses of 2, 10 or 50 µg RH5.1 formulated in AS01_B_ adjuvant at monthly intervals (2-2-2, red, N=12; 10-10-10, blue, N=27; 50-50-50, green, N=9) or a “delayed-fractional regimen” of two doses of 50 µg RH5.1 at 0 and 1 months and a third dose of 10 µg RH5.1 at 6 months (50-50---10, purple, N=12). Individual responses are shown as measured 2-4 weeks post-third vaccination, with boxes indicating minimum, maximum and median. (**C**) Sera from volunteers receiving the 10-10-10 regimen of RH5.1/AS01_B_ (N=15) were diluted 1:100 and tested against linear overlapping peptides spanning the RH5 vaccine insert, colour-coded as per panel (**A**). Median, interquartile range (IQR), and range are shown for each peptide. (**D**) Nine pooled total IgGs from the VAC063 study were tested by GIA with or without the indicated recombinant protein in two (RH5-Nt) or three (RH5.1 and RH5ΔNL) independent assays. The total IgGs were tested in a range from 3 to 9 mg/mL, at which each IgG showed ∼60-70% GIA on average (in the absence of protein). In each assay, % GIA Reversal was calculated as 100 x (1 – % GIA with protein / % GIA without protein), and an average % GIA Reversal from two or three assays in individual IgGs (symbols) are shown with the median (bar) of the nine test IgGs. (**E**) *In vitro* GIA of RH5.1-specific or RH5ΔNL-specific IgG affinity-purified from a pool of human sera collected two weeks post-final vaccination with RH5.1/AS01_B_. The EC_50_ (concentration of antigen-specific polyclonal IgG that gives 50% GIA, dashed line) was calculated by non-linear regression: RH5.1, r^2^=0.98, N=19; RH5ΔNL, r^2^=0.99, N=20).

We next assessed whether IgG antibodies targeting these different structural regions contribute to functional growth inhibition of *P. falciparum* parasites *in vitro* by first using an “antigen-reversal” GIA assay. As expected, inclusion of recombinant RH5.1 protein in the GIA assay could completely reverse all GIA mediated by a pool of purified IgG from RH5.1/AS01_B_ vaccinees. The same result was obtained when using the same concentration of RH5ΔNL protein. In contrast, no reversal of GIA was observed when using recombinant RH5-Nt, even at 8-fold higher molar concentration (**Fig. 1D**). We also affinity-purified anti-RH5.1 and anti-RH5ΔNL human IgG and both samples showed high level growth inhibition. Following titration in the GIA assay, the RH5ΔNL-specific IgG showed an ∼9-fold improvement in terms of the antigen-specific EC_50_ (8 µg/mL, 95% CI: 6-27), as compared to RH5.1-specific IgG (70 µg/mL, 95% CI: 50-114) (**Fig. 1E**). These data show antibodies targeting the N-terminus or intrinsic loop of RH5 do not contribute to functional GIA induced by the RH5.1 vaccine candidate. These data did not assess the small C-terminus of RH5, however, we isolated a novel human IgG mAb, called R5.CT1, from an RH5.1/AS01_B_ vaccinee, that recognized this region. The R5.CT1 clone specifically bound peptides 61 and 62 (**Fig. S1A**) which together span the C-terminal 20 amino acids of RH5. Notably mAb R5.CT1 showed no GIA against *P. falciparum in vitro* (**Fig. S1B**). Together, these data suggest vaccine-induced human anti-RH5.1 IgG growth inhibitory antibodies recognize the alpha-helical core of the RH5 molecule and not the disordered regions. Also, the reason purified polyclonal RH5ΔNL-specific IgG is substantially more potent than RH5.1-specific IgG on a per µg basis is most likely due to the loss of these non-growth inhibitory responses.

### RH5ΔNLC^HS1^-SpyTag vaccine induces similar growth inhibitory antibodies to RH5ΔNL

In light of the above data and to initiate design of an improved RH5-based vaccine candidate, we first assessed three constructs based on the original design of the RH5ΔNL molecule. All three constructs were produced as soluble secreted proteins, using the ExpreS^2^ *Drosophila* S2 stable cell line platform ^28^, and purified by a C-terminal four amino acid C-tag ^29^ – technologies that we previously used to biomanufacture the full-length RH5.1 protein for clinical trials ^20^. Alongside the existing RH5ΔNL protein, we produced two additional molecules that also included removal of the C-terminal 20 amino acids (to make “RH5ΔNLC”) followed by addition of a C-terminal SpyTag (ST), prior to the C-tag, to enable conjugation to SpyCatcher (SC)-based display platforms ^24,26^. The first of these two new molecules otherwise maintained the same RH5 sequence, which we termed “RH5ΔNLC-ST”. The second version, RH5ΔNLC^HS1^-ST, used a previously reported RH5 sequence bearing 18 mutations, defined *in silico*, that confer improved molecular packing, surface polarity and thermostability of the molecule without affecting its ligand binding or immunogenic properties ^30^ (**Fig. 2A**). Each protein was subsequently expressed from a polyclonal S2 stable cell line and purified from the supernatant by C-tag affinity and size exclusion chromatography. Purified proteins ran at their expected molecular weights on an SDS-PAGE gel (**Fig. 2B**). RH5ΔNLC-ST protein was also recognized by a panel of 14 human mAbs previously shown to span six distinct conformational epitope regions on the RH5 molecule ^31^ (**Fig. S2A**). Notably, the RH5ΔNLC^HS1^-ST protein showed greatly reduced or no mAb binding to one of these epitope sites, and loss of binding of a single mAb at another site (**Fig. S2B**), likely due to the introduction of the stabilizing mutations in this variant RH5 construct ^30^. Conversely, an approximately 8-fold higher yield on average of purified RH5ΔNLC^HS1^-ST protein was achieved, as compared to RH5ΔNLC-ST and as anticipated when including the stabilizing mutations (**Fig. 2C**).

**Figure 2.**
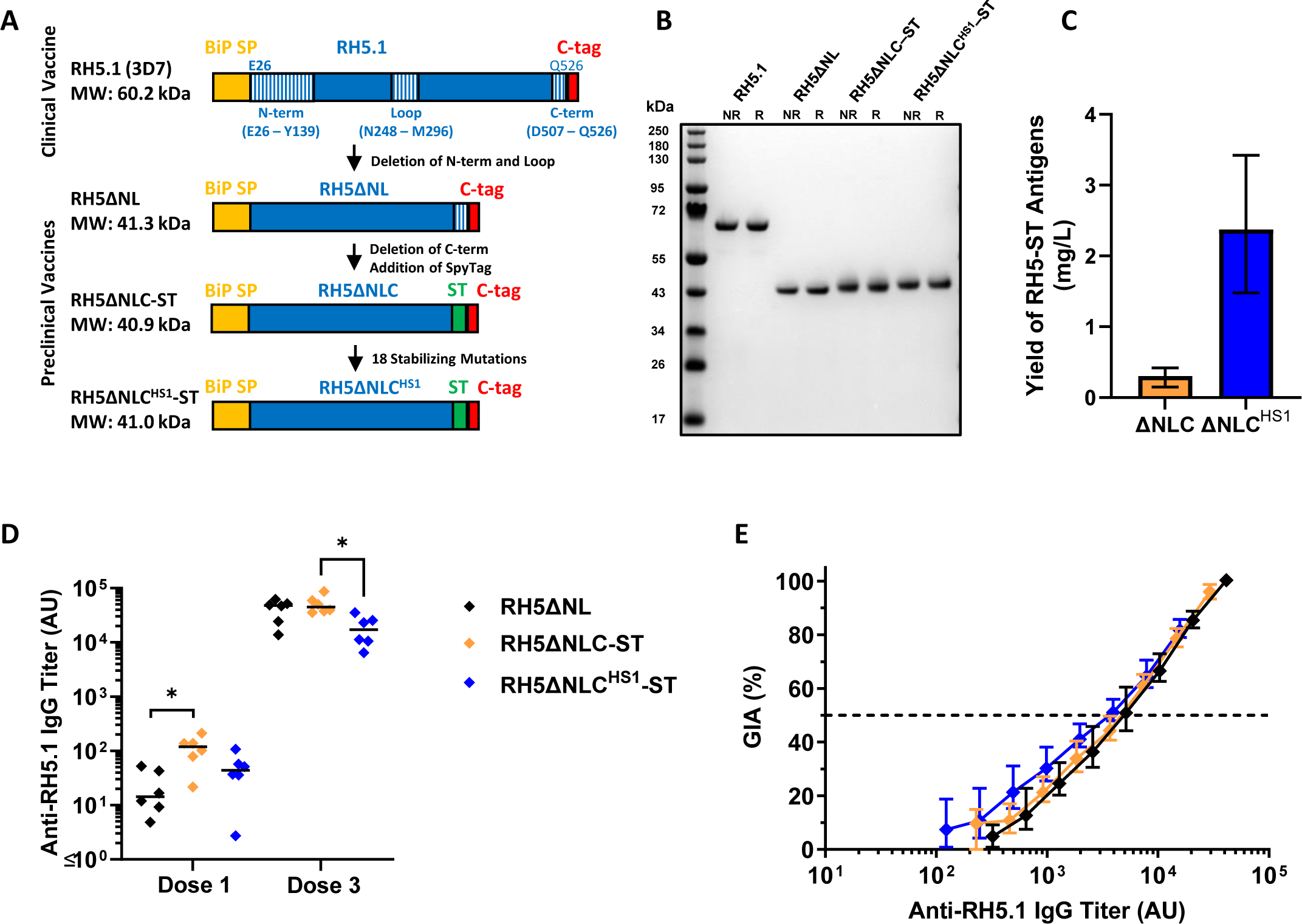
Expression and immunogenicity testing of SpyTagged-RH5ΔNLC constructs. (**A**) RH5 vaccine constructs based on *P. falciparum* 3D7 sequences. All have an N-terminal *Drosophila* BiP secretion signal peptide (SP; which is cleaved off during expression) and end with a C-terminal C-tag for affinity purification. Constructs with a SpyTag (ST) included a flexible (GSG)_3_ linker preceding the ST to facilitate epitope accessibility once conjugated to a VLP bearing SpyCatcher. The predicted molecular weight (MW) of each construct based on the primary sequence and the relevant sequences of RH5 N-term, loop and C-term are shown. (**B**) Non-reduced (NR) and reduced (R) SDS-PAGE gel of affinity and SEC purified RH5.1, RH5ΔNL, RH5ΔNLC-ST and RH5ΔNLC^HS1^-ST proteins. (**C**) Final yield of RH5 protein (in mg) purified from one liter of *Drosophila* S2 stable cell line supernatant. Bars show the mean yield and error bars the range from N=3 independent purification campaigns for each protein. (**D**) BALB/c mice (N=6 per group) were immunized intramuscularly with three 2 µg doses (on days 0, 21 and 42) of RH5ΔNL, RH5ΔNLC-ST or RH5ΔNLC^HS1^-ST, all formulated in Matrix-M™ adjuvant. Anti-RH5 (full-length RH5.1) IgG titers were measured in the serum by ELISA after dose 1 (day 20) and dose 3 (day 70). Each point represents a single mouse and the line the median. Analyses using Kruskal-Wallis test with Dunn’s multiple comparison test across the three groups at each time-point; \**P* < 0.05. (**E**) A single-cycle *in vitro* GIA assay against 3D7 clone *P. falciparum* parasites was performed with total purified IgG from pooled mouse sera (N=6 mice pooled per group). GIA is plotted against the anti-RH5 (full-length RH5.1) titer measured by ELISA in each purified total IgG to assess functional antibody quality, i.e., GIA per unit anti-RH5.1 IgG. Data show titration curve for each sample, with points showing the mean and range of N=3 replicates per test condition.

To assess immunogenicity of the new SpyTagged antigens, 2 µg each protein was formulated in Matrix-M™ adjuvant and used to immunize BALB/c mice intramuscularly three times at three-week intervals. Anti-RH5 serum IgG responses were measured against full-length RH5.1 by ELISA after the first and final vaccinations. Following the first immunization, the RH5ΔNLC-ST protein was significantly more immunogenic than RH5ΔNL (*P* = 0.02, Dunn’s multiple comparison test), however responses equalized for these two proteins after three immunizations. In contrast, RH5ΔNLC^HS1^-ST showed significantly lower responses (∼2-3-fold) after three doses as compared to RH5ΔNLC-ST (**Fig. 2D**). This small reduction in recognition of the RH5.1 protein is likely explained by the introduction of the stabilizing mutations into the RH5ΔNLC^HS1^ construct. To determine if the stabilizing mutations in RH5ΔNLC^HS1^-ST and/or C-terminal truncation in RH5ΔNLC would also affect the functional quality of the growth inhibitory antibody response, we purified the total IgG from pools of mouse sera (6 mice per antigen/group) and tested for *in vitro* GIA against *P. falciparum* (**Fig. 2E**). Here all three proteins could induce an anti-RH5 IgG response with very similar functional quality, i.e., same levels of GIA per unit of anti-RH5 IgG. Consequently, given i) the comparable functional quality of anti-RH5 IgG induced by both SpyTagged proteins and ii) the very low production yield of RH5ΔNLC-ST, we elected to progress the RH5ΔNLC^HS1^-ST protein to further study despite the small reduction in overall immunogenicity, and termed this construct “RH5.2-ST”.

### Production of a RH5.2-HBsAg virus-like particle

To produce a new VLP-based vaccine candidate, we next tested conjugation of the RH5.2-ST protein to a hepatitis B surface antigen particle fused to SpyCatcher (HBsAg-SC) ^26^. We initially conjugated the RH5.2-ST to HBsAg-SC in a 1:1 molar ratio. Following an overnight conjugation reaction, any free unconjugated RH5.2-ST protein was removed by SEC, thereby leaving the conjugated RH5.2-HBsAg VLP product. Analysis by reducing SDS-PAGE showed the expected banding pattern for HBsAg-SC with a dominant monomer band (∼37.0 kDa) as well as multimers (**Fig. 3A**). Following conjugation, a new band corresponding to the RH5.2-HBsAg monomer unit was observed at the expected size of ∼77 kDa along with other bands corresponding to the expected multimers at higher molecular weight. Free unconjugated RH5.2-ST protein was not observed following its removal by SEC, although some unconjugated HBsAg-SC monomer units remained within the VLP preparation. Analysis by densitometry indicated a conjugation efficiency of ∼80%. A study in BALB/c mice was performed next to compare the immunogenicity of RH5.2-ST soluble protein versus the RH5.2-VLP. Dosing of the RH5.2-VLP was adjusted in each case to deliver the same molar amount of RH5.2 antigen as the soluble protein comparator (**Fig. 3B**). Following three immunizations, the RH5.2-VLP formulated in Matrix-M™ adjuvant showed comparable anti-RH5 serum IgG responses across all three doses tested (1, 0.1 and 0.01 µg; *P* = 0.39, Kruskal-Wallis test). In contrast, the same analysis with soluble RH5.2-ST protein showed a clear dose response, with no antibodies detected at the lowest 0.01 µg dose (*P* < 0.0001, Kruskal-Wallis test). When comparing across the same doses of soluble RH5.2-ST versus the RH5.2-VLP, only the 1 µg dose showed comparable immunogenicity, whilst the RH5.2-VLP was significantly more immunogenic at the lower doses (*P* = 0.002 for both the 0.1 and 0.01 µg doses, Dunn’s multiple comparison test). Finally, in the absence of adjuvant, a 1 µg dose of the RH5.2-VLP still induced responses, albeit at a lower level than when using Matrix-M™ adjuvant; in contrast the soluble protein showed negligible immunogenicity (**Fig. 3B**). Analysis of responses following the first and second vaccinations also showed that the 1 and 0.1 µg doses of RH5.2-VLP formulated in Matrix-M™ primed detectable serum antibody responses after only a single immunization and achieved maximal titers after two immunizations. In all cases, the RH5.2-VLP was more immunogenic than the soluble protein (**Fig. S3A,B**). A second experiment was performed using the same total dose of antigen to mirror clinical practice. Here, a 16 ng total protein dose of the RH5.2-VLP was compared to a 16 ng dose of soluble RH5.2-ST or soluble RH5.1 (the current lead clinical antigen); all were formulated in Matrix-M™ adjuvant. Following three immunizations, only the RH5.2-VLP showed high titer anti-RH5 serum IgG responses in contrast to negligible immunogenicity observed with either soluble protein vaccine (**Fig. 3C**). Both experiments confirmed the new RH5.2-VLP is inherently more immunogenic than soluble RH5 protein in mice.

**Figure 3.**
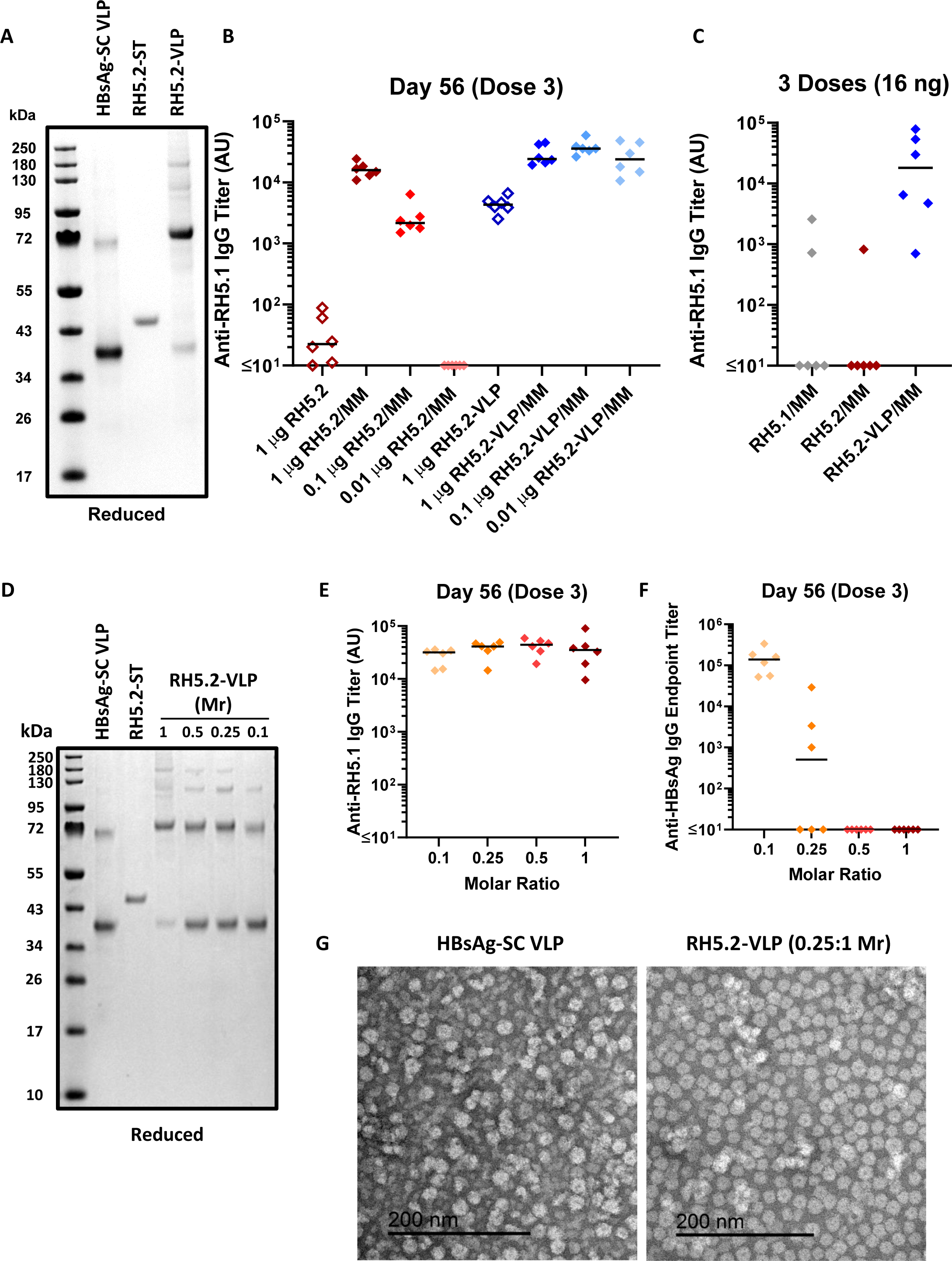
Production and immunogenicity testing of the RH5.2-VLP vaccine candidate. (**A**) Reducing SDS-PAGE gel of HBsAg-SC VLP and RH5.2-ST protein. These proteins were conjugated together in a 1:1 molar ratio. The resulting RH5.2-VLP was SEC purified and is run in the final lane. (**B**) BALB/c mice (N=6 per group) were immunized intramuscularly with three doses of RH5.2-ST protein (“RH5.2”) or RH5.2-VLP on days 0, 21 and 42 either with (closed symbols) or without (open symbols) Matrix-M™ (MM) adjuvant. Dosing of the RH5.2-VLP was adjusted in each case to deliver the same molar amount of RH5.2 antigen as the soluble protein comparator (1, 0.1 or 0.01 µg). Anti-RH5 (full-length RH5.1) IgG titers were measured in the serum by ELISA after three doses at day 56. Each point represents a single mouse and the line the median. (**C**) BALB/c mice (N=6 per group) were immunized intramuscularly with three doses of RH5.1 protein, RH5.2-ST protein (“RH5.2”) or RH5.2-VLP on days 0, 21 and 42. All vaccines used a total dose of 16 ng formulated in Matrix-M™ (MM) adjuvant. Anti-RH5 (full-length RH5.1) IgG titers were measured in the serum by ELISA after three doses at day 56. Each point represents a single mouse and the line the median. (**D**) Reducing SDS-PAGE gel as in panel (**A**) but showing RH5.2-VLP produced by conjugating RH5.2-ST and HBsAg-SC VLP components at the indicated molar ratios (Mr). (**E**) BALB/c mice (N=6 per group) were immunized intramuscularly with three doses of RH5.2-VLP, produced using the indicated molar ratios of RH5.2-ST to HBsAg-SC (0.1:1, 0.25:1, 0.5:1 and 1:1), on days 0, 21 and 42. Dosing was adjusted in each case to deliver the same molar amount of RH5.2 antigen (10 ng); total RH5.2-VLP dose = 232, 52, 40 and 23 ng, respectively. All vaccines were formulated in Matrix-M™ adjuvant. Anti-RH5 (full-length RH5.1) IgG titers and (**F**) anti-HBsAg IgG titers were measured in the serum by ELISA after three doses at day 56. Each point represents a single mouse and the line the median. (**G**) Negatively-stained TEM image of HBsAg-SC VLP starting material and RH5.2-VLP vaccine made using the 0.25:1 molar ratio (Mr). Scale bar 200 nm.

However, despite the highly promising immunogenicity, ongoing studies indicated the conjugated RH5.2-VLP was prone to precipitation during production, resulting in substantial loss of product. We thus attempted to optimize reaction conditions by increasing the salt concentration and lowering the temperature, as well as by combining the two components (RH5.2-ST and HBsAg-SC) dropwise. We also tested incubation of the two components in different molar ratios (RH5.2-ST:HBsAg-SC as 1:1, 0.5:1, 0.25:1 and 0.1:1); here, as expected, more unconjugated HBsAg-SC monomer units remained when combining the VLP with less RH5.2-ST (**Fig. 3D**). Precipitation was also greatly decreased, and overall process yield increased when using the 0.25:1 or 0.1:1 molar ratios in the conjugation reaction. We therefore next proceeded to screen the different products for immunogenicity. BALB/c mice were immunized three times with the four different RH5.2-VLPs all formulated in Matrix-M™ adjuvant.

Dosing was adjusted in each case to deliver the same molar amount of RH5.2 antigen (10 ng). Interestingly, maximal titers were reached faster with VLPs produced using the lower molar ratios (**Fig. S4A,B**), although following three doses all preparations showed comparable anti-RH5.1 serum IgG responses (**Fig. 3E**). Serum antibody responses against the HBsAg VLP carrier inversely related to the molar ratio used in the conjugation reaction, with no detectable responses in mice immunized with the RH5.2-VLP produced using the 1:1 or 0.5:1 ratio (**Fig. 3F**). These higher anti-HBsAg responses, especially in the 0.1:1 molar ratio group, could have been due to the higher total protein dose used in this experiment and/or excess of unconjugated HBsAg-SC subunits on these particles. Nevertheless, we proceeded with further study and evaluation of the RH5.2-VLP produced using the 0.25:1 ratio, as a balanced trade off with regard to production yield versus strong anti-RH5.2 immunogenicity and low anti-HBsAg VLP carrier immunogenicity. Further analysis of this product by transmission electron microscopy (TEM) confirmed particles of the expected ∼20 nm in size (**Fig. 3G**). The RH5.2-VLP was also recognized by the same anti-RH5 human mAbs as reacted with the parental RH5ΔNLC^HS1^-ST protein, confirming the presence and accessibility of these critical conformational epitopes on the VLP (**Fig. S2C**).

### The growth inhibitory antibody response induced by the RH5.2-VLP is superior to RH5.1.

In a final study, we compared the functional immunogenicity of the RH5.2-VLP to soluble RH5.2-ST protein and the current clinical antigen (soluble RH5.1 protein) in Wistar rats. All antigens were formulated in Matrix-M™ adjuvant and administered intramuscularly. Groups of six animals were immunized three times, at monthly intervals, using the same total dose of vaccine (2 µg) to mirror clinical practice. Serum IgG antibody levels were assessed against full-length RH5.1 by ELISA. Responses induced by RH5.1 and the RH5.2-VLP reached maximal levels after two doses, and were superior to soluble RH5.2-ST after every vaccine dose, with RH5.1 significant versus RH5.2-ST (**Fig. 4A**).

**Figure 4.**
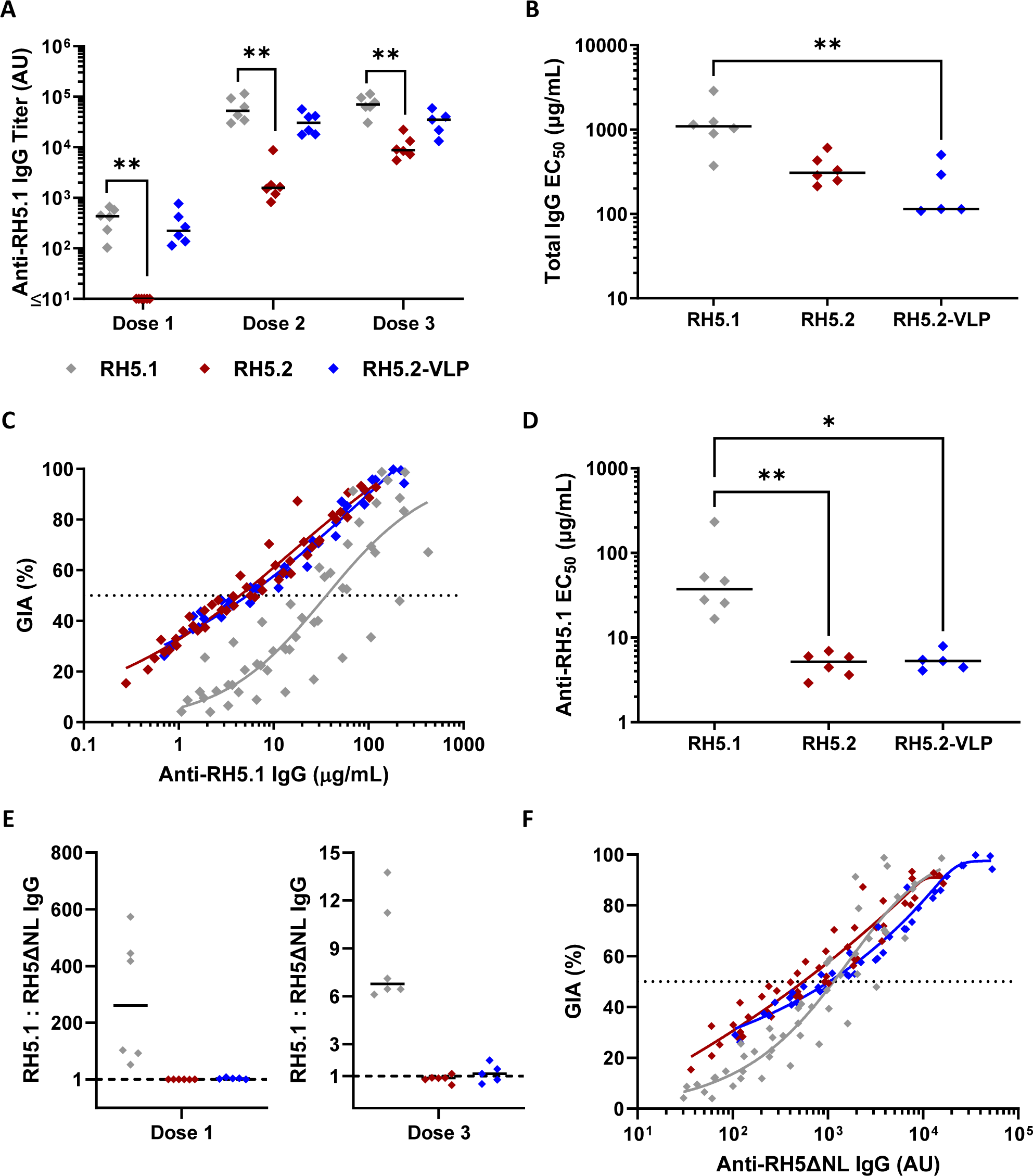
Functional immunogenicity testing of RH5.1, RH5.2 and the RH5.2-VLP in rats. (**A**) Wistar rats (N=6 per group) were immunized intramuscularly with three doses of RH5.1 protein, RH5.2-ST (“RH5.2”) protein or RH5.2-VLP on days 0, 28 and 56. All vaccines used a total dose of 2 µg formulated in Matrix-M™ adjuvant. Anti-RH5 (full-length RH5.1) IgG titers were measured in the serum by ELISA after each dose on days 14, 42 and 70, respectively for Doses 1-3. Each point represents a single rat and the line the median; N=5 for Dose 3 of RH5.2-VLP as a single rat was euthanized after a problem with a study-related procedure. Analysis using Kruskal-Wallis test with Dunn’s multiple comparison test across the three vaccine groups with each Dose result analyzed separately; \*\**P* < 0.01. (**B**) A single-cycle *in vitro* GIA assay against 3D7 clone *P. falciparum* parasites was performed with total IgG purified from serum from each vaccinated rat post-final immunization (N=5-6 per group). Total IgG was titrated in the assay, and the concentration in mg/mL required to achieve 50 % GIA (EC_50_) was interpolated. Data show the EC_50_ for each rat and the line the median. Analysis using Kruskal-Wallis test with Dunn’s multiple comparison test; \*\**P* < 0.01. (**C**) GIA data plotted against the anti-RH5.1 IgG concentration measured by quantitative ELISA in each purified total IgG to assess functional antibody quality, i.e., GIA per µg anti-RH5.1 IgG. A non-linear regression curve is shown for all samples combined in each vaccine group (RH5.1: r^2^=0.75, N=144; RH5.2: r^2^=0.93, N=143; RH5.2-VLP: r^2^=0.96, N=120). The dashed line indicates 50 % GIA. (**D**) The concentration of RH5.1-specific IgG in µg/mL required to achieve 50 % GIA (EC_50_) was interpolated by non-linear regression for each individual rat from the data in (**C**). Data show the EC_50_ for each rat and the line the median. Analysis using Kruskal-Wallis test with Dunn’s multiple comparison test; \**P* < 0.05, \*\**P* < 0.01. (**E**) Ratio of the serum IgG ELISA response as measured using the RH5.1 and RH5ΔNL proteins after the first and third vaccinations. Data shown for each rat (N=5-6 per group) and the line the median. (**F**) GIA data plotted against the anti-RH5ΔNL IgG titer measured by ELISA with arbitrary unit (AU) readout in each purified total IgG to assess functional antibody quality, i.e., GIA per unit anti-RH5ΔNL IgG. A non-linear regression curve is shown for all samples combined in each vaccine group (RH5.1: r^2^=0.86, N=144; RH5.2: r^2^=0.90, N=143; RH5.2-VLP: r^2^=0.95, N=120). The dashed line indicates 50 % GIA.

Following the third dose, total IgG was purified from serum and titrated in the assay of GIA against *P. falciparum* parasites. Here, the RH5.2-VLP showed significantly improved GIA over RH5.1, with the median EC_50_ of total IgG 9.6-fold lower (**Fig. 4B**). Given the comparable quantitative immunogenicity shown by RH5.1 and the RH5.2-VLP (**Fig. 4A**), we next assessed the functional quality of the RH5.1-specific IgG by plotting the GIA data versus ELISA performed on the purified total IgG (**Fig. 4C**). Here, the antibodies induced in the RH5.2-VLP and RH5.2-ST protein immunized groups showed identical quality, i.e., the same GIA per µg of RH5.1-specific IgG, and both significantly improved upon the functional quality induced by RH5.1 immunization (**Fig. 4D**). These data suggest the functional quality of RH5.2-induced IgG is comparable with soluble protein or VLP delivery. Consequently, the improvement in overall levels of GIA observed with the RH5.2-VLP (**Fig. 4B**) relate to its superior quantitative immunogenicity (when comparing to soluble RH5.2) and superior qualitative immunogenicity (when comparing to RH5.1). Given our earlier data indicated the N-terminus and intrinsic loop of RH5 do not contribute to functional GIA induced by RH5.1 in humans, we hypothesized the improvement in functional antibody quality seen with RH5.2 over RH5.1 in the rats was due to loss of responses against these disordered regions of the molecule. We thus tested the rat sera by ELISA against RH5ΔNL and compared the ratio of this response to the RH5.1 response (**Fig. 4E**). As expected, the ratios for RH5.2 and RH5.2-VLP were approximately one, given the RH5.2 immunogen is based on the RH5ΔNL structure, and thus ELISA with either RH5.1 or RH5ΔNL should give a comparable readout. However, the ratio of RH5.1:RH5ΔNL-specific IgG induced by the RH5.1 vaccine was ∼250 following the first dose, suggesting the RH5ΔNL antibody response is initially sub-dominant to responses against the N-terminus and intrinsic loop present in RH5.1. This sub-dominance decreases after three vaccine doses, with the ratio reduced to ∼6.5 (**Fig. 4E**). Overall, these ELISA data suggested a substantial antibody response is mounted to the N-terminus and/or intrinsic loop when using RH5.1. To explore this further, we also conducted a second quality analysis by re-plotting the GIA data versus ELISA on the purified IgG performed against RH5ΔNL protein. Here, in support of our hypothesis, all three constructs now performed similarly, with each on average achieving 50 % GIA at approximately the same level of anti-RH5ΔNL IgG (**Fig. 4F**). These data strongly suggest all of the GIA induced in the rats by RH5.1 was mediated by the subset of IgG that recognize RH5ΔNL, in agreement with the observations in human vaccine responses (**Fig. 1**).

## Discussion

Various vaccine candidates encoding the RH5_FL molecule have been progressed clinically ^18,19,21^, with the most advanced candidate, RH5.1/Matrix-M™ (ClinicalTrials.gov NCT04318002), now entering a Phase 2b efficacy trial in a malaria-endemic country. These vaccines have been developed and prioritized based on evidence that RH5 vaccine-induced *in vivo* growth inhibition (IVGI) of blood-stage *P. falciparum* is antibody-mediated and correlates with *in vitro* GIA in preclinical and human challenge studies ^17,21–23^. However, the quantity of anti-RH5_FL IgG identified as fully protective in *Aotus* monkeys (immunized with protein formulated in Freund’s adjuvant) was high (12) and at a level not yet observed in clinical testing. We therefore sought to develop a new vaccine candidate with improved quantitative and/or qualitative functional anti-RH5 antibody immunogenicity.

To select a vaccine antigen design, we explored functional responses induced in UK adults vaccinated with the RH5.1/AS01_B_ vaccine candidate ^21^. These ELISA data confirmed vaccinees mounted responses against the whole of the RH5 molecule, including the conformational alpha-helical core of RH5 as well as the three linear disordered regions: the long N-terminal region (RH5-Nt), intrinsic loop and small C-terminus. However, a combination of GIA reversal and antibody depletion assays, and analysis of a human mAb to the C-terminus, all indicated that antibodies raised to the three disordered regions are not making a measurable contribution to the overall levels of GIA mediated by anti-RH5.1 IgG. These data are consistent with previous reports showing that murine or human mAbs targeting the N-terminus or intrinsic loop do not inhibit *P. falciparum* growth *in vitro* ^31,32^ and that high-dose passive transfer of a mAb against the intrinsic loop failed to protect *Aotus* monkeys against *P. falciparum* challenge ^22^. In contrast to the results reported here, another study reported that RH5-Nt binds the parasite protein P113 and that vaccination of rabbits with RH5-Nt protein could induce antibodies that mediate modest levels of GIA ^33^; however, we could not identify a similar contribution of anti-RH5-Nt IgG to the GIA induced by human RH5.1 vaccination in our studies. Our data are also consistent with other studies that have since questioned the significance of P113 in merozoite invasion. In particular, one study found anti-P113 antibodies that block the interaction with RH5-Nt are GIA-negative ^34^, whilst another reported P113 plays an important role in maintaining normal architecture of the parasitophorous vacuole membrane within the infected erythrocyte ^35^. Moreover, it has since been reported that RH5-Nt is cleaved off the RH5 molecule within the micronemes of the *P. falciparum* parasite by the aspartic protease plasmepsin X prior to release of RH5 to the merozoite surface ^36^, suggesting it would be an unlikely target of antibodies. In summary, these data strongly suggest these regions of disorder within RH5_FL are not targets of functional IgG. This conclusion is further supported by our data showing the RH5ΔNL protein could reverse all GIA induced by human RH5.1 vaccination, suggesting that all growth inhibitory epitopes targeted by the human IgG are located within this protein construct; this is also consistent with known epitope information of anti-RH5 murine and human mAbs reported previously and shown to have anti-parasitic activity ^27,31,32,37^. Finally, our data showed an ∼9-fold improvement in the antigen-specific GIA EC_50_ potency when comparing affinity-purified polyclonal IgG specific for RH5ΔNL versus RH5.1. On the basis of all these data, we elected to focus new vaccine design efforts on a molecule lacking all three disordered regions, which we termed “RH5ΔNLC”.

To prepare for biomanufacture of clinical-grade immunogen, we produced new protein constructs utilizing the same expression and purification platform technologies as those used previously for the clinical biomanufacture of RH5.1 ^20^. Here we could produce the wild-type RH5ΔNLC antigen with a C-terminal SpyTag followed by C-tag, as well as a second variant sequence incorporating 18 mutations that were defined *in silico* and previously reported to confer improved molecular packing, surface polarity and thermostability of the RH5ΔNL molecule ^30^. Consistent with the original report, the yield of purified SpyTagged RH5ΔNLC protein was ∼8-fold higher when incorporating the stabilizing mutations.

Immunization of mice with these monomeric soluble proteins-in-adjuvant showed a modestly higher antibody response, as measured against RH5.1 antigen, when using the wild-type sequence proteins as compared to the mutated version. Consistent with this, we noted reduced binding or loss of binding by human mAbs at two previously identified antigenic sites within the RH5ΔNL molecule ^31^, indicating that a small number of specific antibody epitopes were affected by the stabilizing mutations. However, regardless of this and consistent with the original report of the stabilized RH5ΔNL sequence ^30^, we observed no difference in the functional quality of the RH5-specific antibodies elicited through vaccination of mice with these new SpyTagged proteins formulated in Matrix-M™ adjuvant. Whether these mutations would significantly impact immune responses in humans remains to be determined. Given the substantially higher yield of the stabilized RH5ΔNLC SpyTagged variant, we proceeded with this stabilized version of the RH5ΔNLC protein which we termed “RH5.2-ST”.

VLP-based immunogens, that deliver multimeric or arrayed antigen, have been widely shown to offer numerous advantages over soluble antigen vaccines. These include improved trafficking to draining lymph nodes, improved efficiency of B cell receptor cross-linking as well as oriented antigen display, all of which can substantially improve quantitative and/or qualitative antibody immunogenicity ^38,39^. We thus explored the delivery of the RH5.2 immunogen following bioconjugation to HBsAg VLPs using the SpyTag-SpyCatcher platform ^24,26^. These lipoprotein VLPs are ∼20-30 nm in size and contain ∼100 monomeric HBsAg polypeptide subunits ^40^. We selected HBsAg VLPs given they have been safely used in humans for decades as a highly effective anti-hepatitis B virus vaccine ^41^, and to align RH5.2 delivery platform with the two approved pre-erythrocytic malaria vaccines, RTS,S/AS01 (Mosquirix™) and R21/Matrix-M™, both of which are adjuvanted chimeric HBsAg VLPs. Our initial attempts to conjugate RH5.2-ST to HBsAg-SC VLPs at a 1:1 molar ratio showed a maximal conjugation efficiency of ∼80% by densitometry analysis, however, the overall process yield was low due to significant precipitation and loss of product during the conjugation process. Reaction conditions were subsequently optimized leading to improved yield, but this necessitated conjugating RH5.2-ST at a lower molar ratio. Efforts to re-design RH5-based protein immunogens with improved solubility characteristics currently remain the focus of ongoing work.

We subsequently undertook a series of mouse immunogenicity studies comparing the RH5.2-VLP versus the soluble RH5.2 and RH5.1 vaccine candidates. Notably, quantitative antibody immunogenicity in mice was determined by the presence of adjuvant, number of immunizations and immunogen dose. These data showed the RH5.2-VLP was consistently more immunogenic than soluble antigen after three immunizations and when tested i) at low dose (in the 10-100 ng range) in the presence of Matrix-M™ adjuvant and ii) at high dose (1 µg) in the absence of Matrix-M™ adjuvant. Responses induced by the RH5.2-VLP in Matrix-M™ adjuvant were also higher after one or two immunizations and reached maximal titers earlier, across the dose range tested, as compared to soluble antigen. Moreover, similar to a previous study of the transmission-blocking malaria antigen Pfs25 conjugated to HBsAg VLPs ^26^, maximal anti-RH5 serum IgG responses were achieved following three immunizations of the RH5.2-VLP in Matrix-M™ adjuvant, regardless of RH5.2 conjugation density to the HBsAg VLP.

We subsequently proceeded to further test the RH5.2-VLP produced using the 0.25:1 molar ratio. These VLPs were of the expected size and bound the same panel of human anti-RH5 mAbs as the soluble RH5.2 antigen. Immunization of Wistar rats with three 2 µg doses of antigen formulated in Matrix-M™ adjuvant showed the RH5.2-VLP outperformed soluble RH5.2 in terms of quantitative immunogenicity but maintained comparability to the larger soluble RH5.1 protein. However, functional testing of the purified total IgG from RH5.2-VLP vaccinated rats showed significantly improved GIA over RH5.1, with a median 9.6-fold reduction in the EC_50_ of total IgG. To our knowledge, this is the first vaccine candidate to significantly outperform RH5.1 in terms of functional immunogenicity in a preclinical model, including comparison to vaccines targeting the wider RH5 invasion complex ^42^. This improvement was driven by a significantly lower antigen-specific EC_50_ of the vaccine-induced IgG, with RH5.2-vaccinated rats achieving 50 % GIA at median levels of ∼5 µg/mL RH5.1-specific antibody. Notably, this qualitative improvement was consistent across all RH5.2-vaccinated rats, whether immunized with soluble antigen or the HBsAg-VLP. This indicates no added benefit of VLP delivery with regard to qualitative immunogenicity and that the conjugated RH5.2 is likely fully exposed and/or flexibly displayed on the VLP surface. Consistent with this, the mAb panel analysis detected the identical range of epitopes on both soluble and VLP-conjugated RH5.2 antigen, including those in the C-terminal region of RH5.2 ^31,43^ that would be expected to be closer to the VLP surface. In summary, vaccination with the RH5.2 immunogen, itself based on the RH5ΔNLC molecule, induced a serum antibody response of superior functional quality per unit of anti-RH5.1 IgG as compared to the current clinical lead vaccine RH5.1/Matrix-M™. This was most likely due to loss of non-functional IgG responses against the disordered regions of the full-length RH5 molecule (when using RH5.2), which appear to dilute the subdominant and functional IgG induced against the helical core (when using RH5.1). In parallel, VLP-based delivery improved quantitative immunogenicity against the smaller RH5.2 immunogen, thereby leading to the highest levels of GIA observed in rats with the RH5.2-VLP.

The RH5.2-VLP antigen has since completed biomanufacture in line with current good manufacturing practice (cGMP) and is entering Phase 1a/b clinical trials in the United Kingdom and The Gambia (ClinicalTrials.gov NCT05978037 and NCT05357560) formulated in Matrix-M™ adjuvant. These will enable the comparison in humans of the RH5-based immunogens delivered as a soluble protein versus an array on HBsAg-VLPs.

## Methods

### Model of RH5.1

AlphaFold model AF-Q8IFM5-F1 ^44,45^ was imported into ChimeraX software ^46^ version 1.6.1 for visualization of the different structural regions of RH5.1.

### Clinical serum samples

All human serum samples were from the VAC063 clinical trial ^21^. Malaria-naïve healthy UK adult volunteers received three intramuscular doses of the RH5.1 antigen ^20^ formulated in 0.5 mL AS01_B_ adjuvant (GSK) in various dosing regimens as previously described. Serum samples taken two weeks after the second dose or the third and final dose were used in the studies reported here. VAC063 received ethical approval from the UK NHS Research Ethics Service (Oxfordshire Research Ethics Committee A, Ref 16/SC/0345) and was approved by the UK Medicines and Healthcare products Regulatory Agency (Ref 21584/0362/001-0011). Volunteers signed written consent forms and consent was verified before each vaccination. The trial was registered on ClinicalTrials.gov (NCT02927145) and was conducted according to the principles of the current revision of the Declaration of Helsinki 2008 and in full conformity with the ICH guidelines for Good Clinical Practice (GCP).

### Generation of polyclonal Schneider 2 (S2) stable cell lines

All RH5 constructs were based on the *P. falciparum* 3D7 clone sequence and potential N-linked glycosylation sequons were mutated from N-X-S/T to N-X-A. Production of stable S2 cell lines expressing the full-length RH5.1 (residues E26-Q526) and RH5ΔNL (residues K140-K247 and N297-N526) proteins has been described previously ^20,27^. Synthetic genes encoding RH5ΔNLC-ST (residues K140-K247 and N297-N506) or RH5.2-ST (residues K140-K247 and N297-N506 with 18 stabilizing mutations ^30^: I157L, D183E, A233K, M304F, K312N, L314F, K316N, M330N, S370A, S381N, T384K, L392K, T395N, N398E, R458K, N463K, S467A, F505L) were codon optimized for expression in

*Drosophila melanogaster* and included flanking 5’ EcoRI and 3’ NotI sites that were used to subclone each gene into the pExpreS^2^-2 plasmid (ExpreS^2^ion Biotechnologies, Denmark) ^28^. These two SpyTagged RH5 constructs also included an N-terminal BiP insect signal peptide and a C-terminal flexible linker (GSGGSGGSG) followed by SpyTag (AHIVMVDAYKPTK) and C-tag (EPEA) ^24,28,29^. Stable polyclonal S2 insect cells lines were generated through transient transfection with ExpreS^2^TR reagent (Expression Systems) mixed with the relevant plasmid and subsequent culturing under selection with G418 (Gibco) supplemented EX-CELL 420 serum-free media (Merck).

### Expression and purification of recombinant RH5 proteins

Stable monoclonal (RH5.1) or polyclonal (RH5ΔNL, RH5ΔNLC-ST, RH5.2-ST) S2 cells lines were cultured in EX-CELL 420 serum-free media (Merck) supplemented with 100 U/mL penicillin and 100 µg/mL streptomycin (Gibco) at 25 °C and 125 rpm. Cell cultures were scaled up to 2.5 L and the supernatant was harvested 3 days later by centrifugation at 3,250 x*g* for 20 min followed by filtration through a 0.22 µm Steritop™ filter unit. Cell supernatant was then concentrated by Tangential Flow Filtration with a Pellicon 3 Ultracel 10 kDa membrane (Merck Millipore) and loaded onto a 10 mL CaptureSelect™ C-tagXL affinity column that had been equilibrated in Tris-buffered saline (TBS; 20 mM Tris-HCl pH 7.4, 150 mM NaCl). The column was then washed with 10 column volumes (CV) of TBS and protein eluted in 2 M MgCl_2_ supplemented with 20 mM Tris-HCl pH 7.4. Eluted protein fractions were then pooled, concentrated and purified into TBS by size exclusion chromatography (SEC) using a HiLoad 16/600 Superdex 75 or 200 pg column (Cytiva) and an ÄKTA Pure™ Protein Purification System (Cytiva).

The RH5-Nt protein encoded residues F25-K140 followed by rat CD4 domains 3 and 4, a biotin acceptor peptide and a C-terminal hexa-histidine tag ^33^. The protein was expressed in Expi293 cells and purified by immobilized metal affinity chromatography (IMAC) using Ni^2+^ resin followed by SEC, with protein eluted into TBS as previously described ^33^.

### SDS-PAGE

Samples were prepared in 1 x Laemmli buffer with or without 1 x dithiothreitol (Biorad). Samples were then heated for 10 min at 95 °C and loaded onto a precast NuPAGE™ 4-12 % Bis-Tris polyacrylamide gel in NuPAGE™ MES SDS running buffer (Thermo Fisher Scientific). Electrophoresis was performed at 200 V for 45 min and gels were stained overnight with Quick Coomassie stain (Protein Ark), destained in distilled water and imaged using an iBright™ FL1500 Imaging System (Thermo Fisher Scientific).

### Production of anti-RH5 monoclonal antibodies

The isolation, expression and purification of the human anti-RH5 monoclonal antibodies (mAbs) used here has previously been described ^31^. In brief, anti-RH5 mAbs were expressed by transient transfection of Expi293 cells with the heavy and light chain plasmids at a 1:1 ratio (0.5 µg of each plasmid per mL of culture). The supernatant was harvested 5-7 days later by centrifugation at 3,250 x*g* for 20 min, filtered through a 0.22 µm filter and then loaded onto a 5 mL Protein G HP column equilibrated in TBS. The Protein G column was washed with 10 CV of TBS and mAbs were eluted in 0.1 M glycine pH 2.7 and neutralized with Tris-HCl pH 9.0. Eluted mAbs were then buffer exchanged into TBS pH 7.4 using 30 kDa Amicon Ultra-15 centrifugal filters (Millipore).

### Conjugation of RH5.2-ST to HBsAg-SC VLPs

Design, expression and purification of HBsAg VLPs with an N-terminal SpyCatcher moiety on each monomer unit (HBsAg-SC) have been previously reported in detail ^26^. Soluble RH5.2-ST protein and HBsAg-SC VLPs were thawed on ice and supplemented with 200 mM NaCl. While on ice, 0.01-0.1 M of RH5.2-ST was added every 10 min to a fixed amount of HBsAg-SC until a final molar ratio of 1, 0.5, 0.25 or 0.1 of RH5.2-ST antigen to HBsAg-SC VLP was achieved; the reaction was then incubated overnight at 4 °C. Conjugation reactions were then loaded onto a Superdex 200 10/300 Increase or Superose 6 10/300 GL Increase SEC column (Cytiva) and purified into 20 mM Tris-HCl pH 7.4, 350 mM NaCl. The SEC purification removed any free excess RH5.2-ST protein, thereby leaving the purified conjugated RH5.2-VLPs (here each VLP is now composed of a mixture of monomer units of RH5.2-ST-SC-HBsAg, i.e., those monomer units onto which the RH5.2-ST had conjugated, and also excess HBsAg-SC monomer units onto which no RH5.2-ST had conjugated). The protein concentration of the purified VLPs was measured using a Pierce™ BCA Protein Assay kit (Thermo Fisher). VLPs were then flash frozen in liquid nitrogen and stored at −80 °C until use. Conjugation reactions were run on SDS-PAGE, and conjugation efficiency (% of HBsAg-SC monomer units in the VLP conjugated to RH5.2-ST) was assessed by densitometry.

### Negative staining transmission electron microscopy (TEM)

VLPs, at 0.1 mg/mL test concentration, were adsorbed onto 200 mesh formvar/carbon copper grids for 1-2 min, washed with Milli-Q water and blotted with filter paper. Grids were then stained with 2 % uranyl acetate for 10-30 s, air dried and imaged using a FEI Tecnai T12 transmission electron microscope.

### Rodent immunization studies

All mouse experiments and procedures were performed under the UK Animals (Scientific Procedures) Act Project Licence (PPL PA7D20B85) and were approved by the University of Oxford Animal Welfare and Ethical Review Body. Eight-week-old female BALB/c mice (Envigo RMS, UK) (N = 5-6 per group) were immunized intramuscularly (i.m.) with 5 µg Matrix-M™ adjuvant (Novavax) alone or 0.01-2 µg test antigen formulated with Matrix-M™ adjuvant on days 0, 21 and 42. Serum was harvested from blood collected from mouse tail veins on day 20, day 41 and by cardiac puncture on day 56. Serum was then stored at −80 °C.

The rat immunization study was performed at Noble Life Sciences, Inc (Maryland, USA). Female Wistar IGS rats (N=6 per group) between 150-200 g (8-12 weeks old) were immunized i.m. with 2 µg antigen formulated in 25 µg Matrix-M™ adjuvant (Novavax) on days 0, 28 and 56. Serum was harvested from the blood following retro-orbital bleeding on days −2, 14, 42 and cardiac puncture on day 70. Serum samples were then frozen and shipped to the University of Oxford, UK for testing.

### Monoclonal antibodies

The R5.CT1 mAb was isolated from a single IgG^+^ memory B cell in the peripheral blood mononuclear cells of an RH5.1/AS01_B_ human vaccinee using a full-length RH5 probe and methodology as described in detail elsewhere ^47^; The antibody genes were cloned into vectors encoding the human IgG1 backbone for expression in HEK293F cells followed by purification. Production of the 2AC7, R5.016 and EBL040 mAbs has been described previously ^31,32,48^.

### Monoclonal antibody ELISA

96-well flat-bottom NUNC Maxisorp plates were coated with 50 µL (2 µg/mL of antigen) RH5ΔNLC-ST, RH5.2-ST or RH5.2-VLP overnight at 4 °C. Plates were washed five times with PBS/Tween-20 (0.05% v/v; PBS/T) and blocked with 200 µL Blocker™ Casein in PBS (Thermo Fisher Scientific) for 1 h at RT. The anti-RH5 human IgG1 mAbs used in this study have been reported previously ^31^. An irrelevant human IgG1 mAb was used as a negative control. Test mAbs were added in triplicate wells at 1 µg/mL (50 µL/well) and plates were incubated at RT for 1 h, washed in PBS/T and then incubated with 50 µL γ-chain specific goat anti-human IgG-alkaline phosphatase (AP) (Thermo Fisher) at a 1/2000 dilution for 1 h at RT. Plates were washed, then developed with 100 µL *p*-nitrophenylphosphate (pNPP) (Thermo Fisher Scientific) substrate in 1 x diethanolamine buffer, read at 405 nm on an ELx800 absorbance microplate reader (Biotek) and analysed with Gen5 software v3.11.

### Standardized ELISAs

Mouse, rat or human anti-RH5.1, -RH5ΔNL or -RH5Nt IgG ELISAs were performed on serum or purified IgG samples using a standardized methodology, as previously described ^18,49^. In brief, plates were coated with 2 µg/mL test antigen in PBS overnight at 4 °C, washed in PBS/T and blocked for 1 h at RT with 200 µL StartingBlock™ or Blocker™ Casein in PBS (Thermo Fisher Scientific). Serum or purified IgG samples were diluted in blocking buffer, added to the plate and incubated for 1 h at RT, prior to washing and incubation with a goat anti-mouse, -rat or -human-IgG-AP secondary antibody (1:2000) for 1 h. Plates were then developed as per the mAb ELISA. Arbitrary units (AU) were assigned to the reciprocal dilution of the standard curve at which an optical density (OD) of 1 was observed. Using Gen5 ELISA software v3.11 the standard curve was used to assign AU to test samples and where possible, calibration-free concentration analysis (CFCA) was used to convert these values into µg/mL ^18,50^.

### RH5 peptide ELISAs

Methodology for ELISA using biotinylated 20-mer peptides overlapping by 12 amino acids covering the full-length RH5 sequence was reported in detail previously ^18^. RH5.1 and RH5-Nt protein (at 2 µg/mL) were adsorbed to 96-well NUNC-Immuno Maxisorp plates (Thermo Fisher Scientific) and test peptides (at 10 µg/mL) were adsorbed to streptavidin plates (Pierce) overnight at 4 °C. Test purified human IgG samples and a negative pre-immunization control IgG, from VAC063 trial vaccinees ^21^, were normalized to 100 µg/mL in Blocker™ Casein in PBS (Thermo Fisher Scientific) and added to triplicate wells following blocking with Blocker™ Casein in PBS. Blank test wells used blocking buffer only.

Antibodies were detected using goat anti-human IgG-AP (Sigma) and developed and analysed as per the mAb ELISA. The R5.CT1 human mAb was tested in the same assay at 2 µg/mL concentration.

### Anti-HBsAg endpoint ELISA

96-well flat-bottom NUNC Maxisorp plates were coated overnight at 4 °C with 0.5 mg/mL recombinant HBsAg (BIO-RAD). Plates were washed with PBS/T, blocked in 5 % skimmed milk and test serum samples were added in duplicate wells and diluted down the plate in a two-fold dilution series. Following a 1 h incubation, plates were washed in PBS/T and incubated for 1 h with goat anti-mouse IgG-AP (Merck). Plates were then developed as per the mAb ELISA. Endpoint titers were calculated by determining the point at which the dilution curve intercepts the x-axis at an absorbance value 3 standard deviations greater than the OD for a naïve mouse serum sample.

### Assay of growth inhibition activity (GIA)

GIA assays were performed according to standardized methodology from the GIA Reference Centre, NIAID/NIH, as previously described ^51^. In brief, total IgG was purified from serum using a 5mL HiTrap Protein-G HP (Cytiva) column and antigen-specific IgG was purified using RH5.1 or RH5ΔNL coated resin ^52,53^. All samples were heat inactivated, depleted of anti-erythrocyte specific antibodies, buffer exchanged into RPMI-1640 media and filter sterilized prior to being incubated at varying concentrations with O+ erythrocytes and synchronized *P. falciparum* 3D7 clone trophozoites for 42 h at 37 °C (“one-cycle GIA”). All samples were tested in a two-fold dilution curve starting at a concentration of 5 mg/mL and the final parasitemia was then quantified through biochemical detection of lactate dehydrogenase in order to calculate % GIA. For the antigen reversal GIA assay, test antibodies were pre-incubated with the indicated concentration of recombinant protein, which were dialyzed against RPMI-1640, in a 96-well plate for 45 min at RT followed by a 15 min incubation at 37 °C. Then, trophozoite parasites were added to the plate to start the GIA assay as described above.

### Statistical analysis

All data were analyzed using GraphPad Prism version 10.0.3 for Windows (GraphPad Software Inc., California, USA). All tests used were two-tailed and are described in the text. To analyze the GIA EC_50_ an asymmetric logistic dose-response curve was fitted to GIA titration data with no constraints, and EC_50_ values were interpolated. To compare ELISA or EC_50_ values across different groups of immunized mice or rats a Kruskal-Wallis test with Dunn’s multiple comparison test was performed. A value of *P* < 0.05 was considered significant.

## Author Contributions

Conceived and performed experiments and/or analysed the data: LDWK, DP, JRB, HD, DQ, AML, SES, DJP, AD, BGW, KMc, AR, CAR, VS, JS, CR-S, RAD, ASI, YZ, GG, JJ, YL, KMi, SJD. Performed project management: ARN, RSM, CRK, AJB, LAS, RA, KS. Contributed reagents, materials, and analysis tools: CC, AMM, IC, SJF, CAL, MH, SB. Wrote the paper: LDWK, SJD.

## Acknowledgments

The authors are grateful for the assistance of Julie Furze, Penelope Lane, Fay Nugent, Wendy Crocker, Charlotte Hague, Daniel Alanine, Robert Ragotte, Darren Leneghan, Geneviève Labbé, Carolyn Nielsen, Martino Bardelli, Jenny Bryant, Lana Strmecki and Matt Higgins (University of Oxford); Sally Pelling-Deeves for arranging contracts (University of Oxford); Colleen Woods (PATH-MVI); Jenny Reimer (Novavax); Ken Tucker, Timothy Phares, Jayne Christen, Cecille Browne and Vin Kotraiah (Leidos); Robin Miller (USAID); and all the VAC063 trial participants.

This work was funded in part by the UK Medical Research Council (MRC) [MR/P001351/1] – this UK funded award was part of the EDCTP2 programme supported by the European Union; the European Union’s Horizon 2020 research and innovation programme under a grant agreement for OptiMalVax (733273); the PATH Malaria Vaccine Initiative; the UK MRC Confidence in Concept (CiC) Tropical Infectious Disease Consortium [MC_PC_15040]; and the Wellcome Trust through a Translation Award [205981/Z/17/Z]. This work, as well as the VAC063 clinical trial, was made possible in part through support provided by the Infectious Disease Division, Bureau for Global Health, United States Agency for International Development (USAID), under the terms of the Malaria Vaccine Development Program (MVDP) (AID-OAA-C-15-00071) for which Leidos Inc. was the prime contractor, and under the terms of GH-BAA-2018-Addendum03 (7200AA20C00017), for which PATH is the prime contractor. The GIA assays were supported in part by the Division of Intramural Research of the National Institute of Allergy and Infectious Diseases (NIAID), National Institutes of Health (NIH), and by an Interagency Agreement (AID-GH-T-15-00001) between the USAID MVDP and NIAID, NIH. The findings and conclusions are those of the authors and do not necessarily represent the official position of USAID. This work was also supported in part by the National Institute for Health Research (NIHR) Oxford Biomedical Research Centre (BRC) and NHS Blood & Transplant (NHSBT; who provided material), the views expressed are those of the authors and not necessarily those of the NIHR or the Department of Health and Social Care or NHSBT. GSK had the opportunity to review the manuscript but content is the sole responsibility of the authors. JS held a Wellcome/African Academy of Sciences DELTAS Africa Grant Master’s Studentship [DEL-15-007: Awandare]. BGW held a UK MRC PhD Studentship [MR/N013468/1]. SB and SJD are Jenner Investigators and SJD held a Wellcome Trust Senior Fellowship [106917/Z/15/Z].

## Data and Materials Availability

Requests for materials should be addressed to the corresponding author.

