## Supplemental Information for "Preclinical Development of a Stabilized RH5 Virus-Like Particle Vaccine that Induces Improved Anti-Malarial Antibodies"

### Supplementary Figures

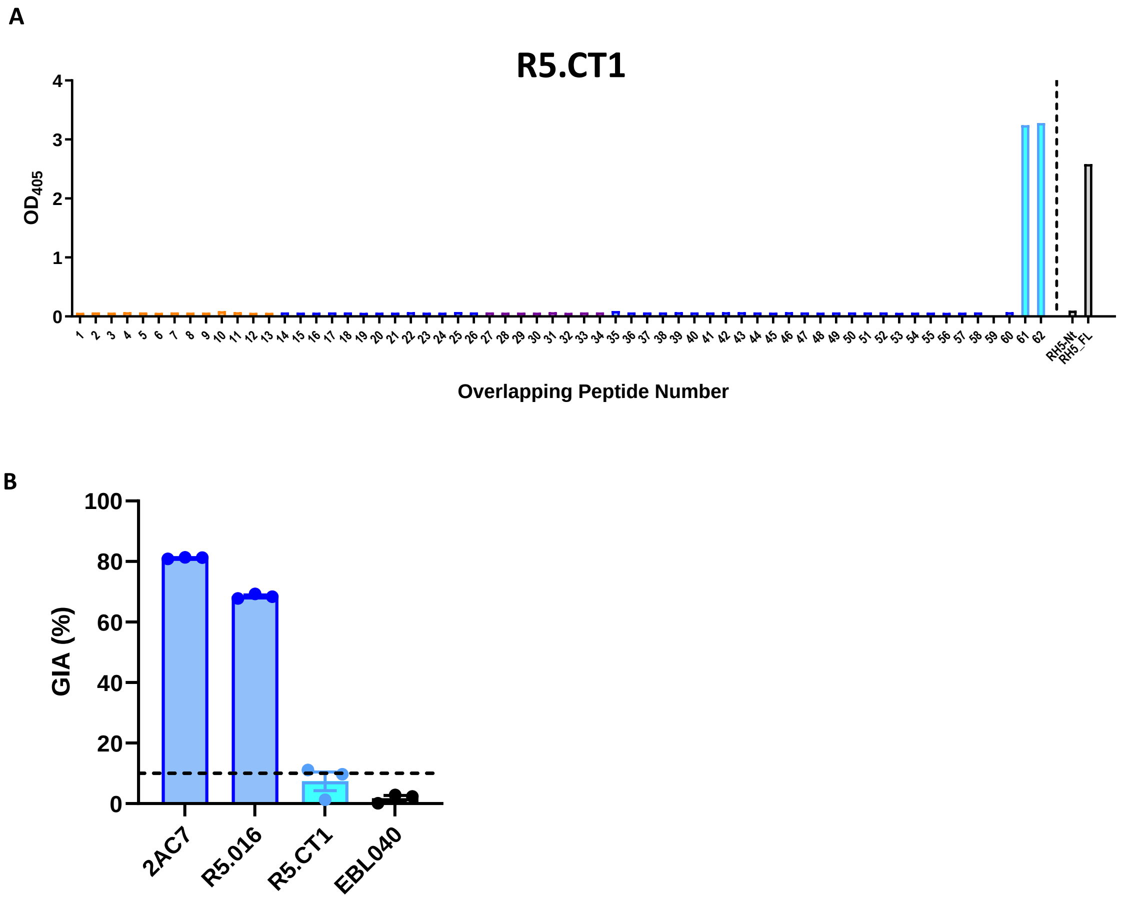


**Figure S1. Assessment of an anti-RH5 C-terminal human mAb.**

(**A**) The recombinant human IgG1 mAb, R5.CT1, was tested by ELISA at 2 µg/mL against linear overlapping peptides spanning the RH5 vaccine insert, colour-coded as per **Figure 1**. Data from single wells are shown, but data are representative of N=3 repeats. Peptides 61 and 62 span the C-terminal 20 amino acids of RH5 and differ by only one amino acid [1]. RH5-Nt and RH5_FL = recombinant protein controls for RH5 N-terminus and full-length, respectively. (**B**) Individual mAbs were tested in triplicate in the GIA assay against 3D7 clone *P. falciparum* parasites. Individual and mean ± SEM GIA % are shown for each mAb. 2AC7 and R5.016 (positive control mAbs) bind RH5ΔNL [2,3] and were tested at 15-20 µg/mL; EBL040 (negative control mAb against Ebola virus) [4] and R5.CT1 were tested at 0.5 mg/mL. Dashed line at 10 % GIA represents typical cut-off for positivity in the assay.

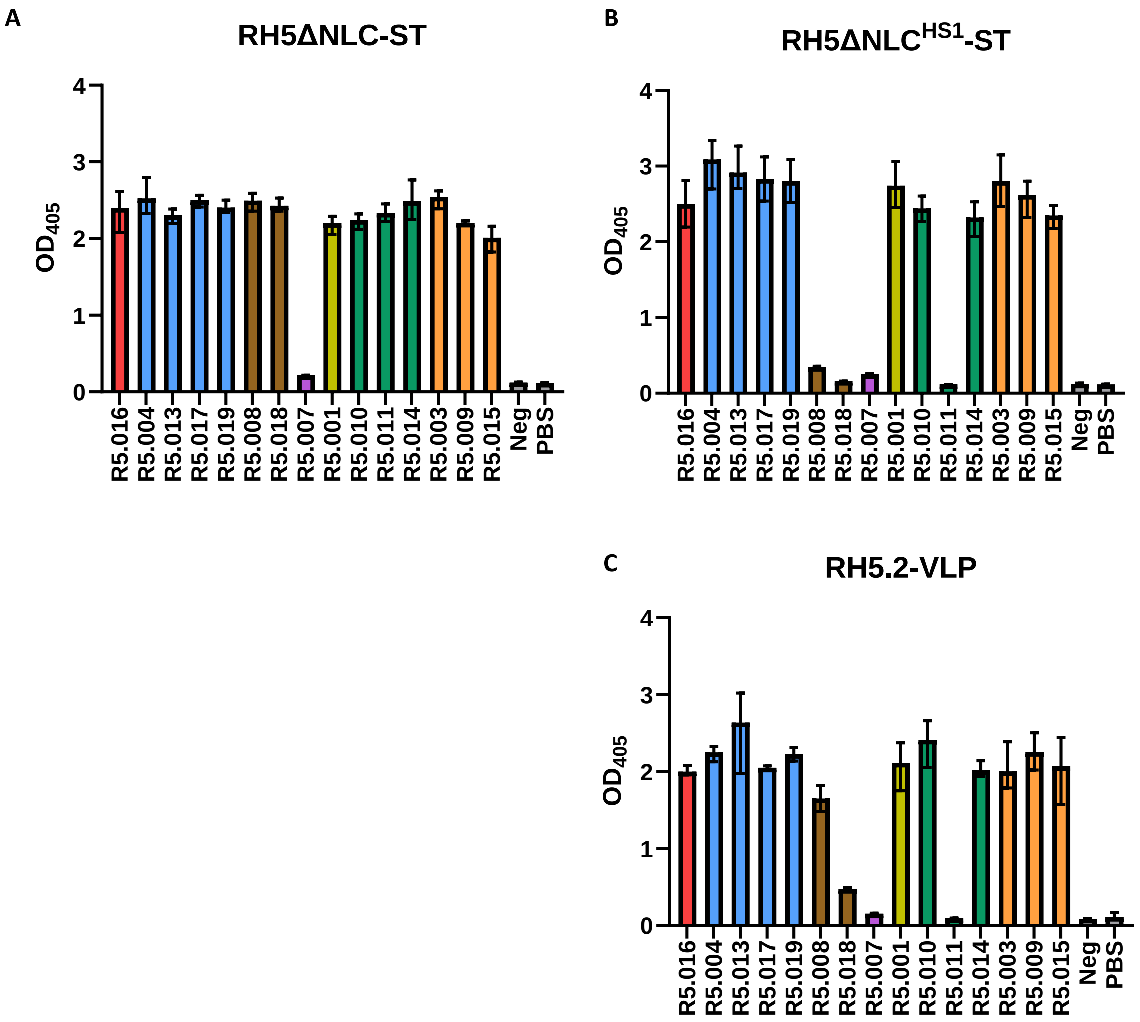


**Figure S2. ELISA to screen for RH5 protein binding to a panel of anti-RH5 human mAbs.**

A binding ELISA was performed on (**A**) RH5ΔNLC-ST protein, (**B**) RH5ΔNLC$^{HS1}$-ST protein and (**C**) RH5.2-VLP using a panel of anti-RH5 human mAbs. This mAb panel is color-coded as previously reported and defines seven epitope regions or antibody competition binding groups across the RH5 molecule [2]. Antibodies of the same color compete for binding, but do not compete with antibodies in other color-coded groups. Clone R5.007 (purple) binds a linear peptide epitope in the intrinsic loop [2] and therefore should not bind to either of these proteins given they lack this sequence. The remaining six groups bind conformational epitopes [2]. The red, blue and brown groups include growth inhibitory antibodies that bind close to or within the basigin binding site on RH5 [2]; the green antibodies do not inhibit invasion but can synergize with other growth inhibitory antibodies [2]; the yellow and orange antibodies do not inhibit parasite growth *in vitro* but block RH5 binding to CyRPA [2,5]. Neg is an irrelevant human IgG1 antibody control. PBS = phosphate-buffered saline only control. Results show the mean and range of optical density at 405 nm ($OD_{405}$) of triplicate wells.

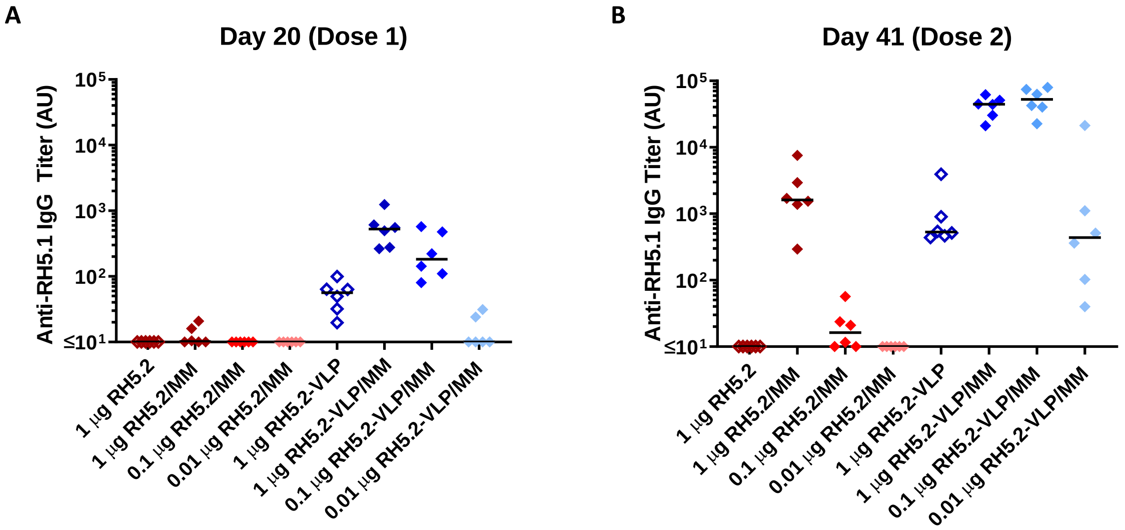


**Figure S3. Immunogenicity testing of the RH5.2-VLP vaccine candidate.**

BALB/c mice (N=6 per group) were immunized intramuscularly with three doses of RH5.2-ST protein or RH5.2-VLP on days 0, 21 and 42 either with (closed symbols) or without (open symbols) Matrix-M™ (MM) adjuvant. Dosing of the RH5.2-VLP was adjusted in each case to deliver the same molar amount of RH5.2 antigen as the soluble protein comparator (1, 0.1 or 0.01 µg). Anti-RH5 (full-length RH5.1) IgG titers were measured in the serum by ELISA after (**A**) dose 1 at day 20, and (**B**) dose 2 at day 41. Each point represents a single mouse and the line the median.

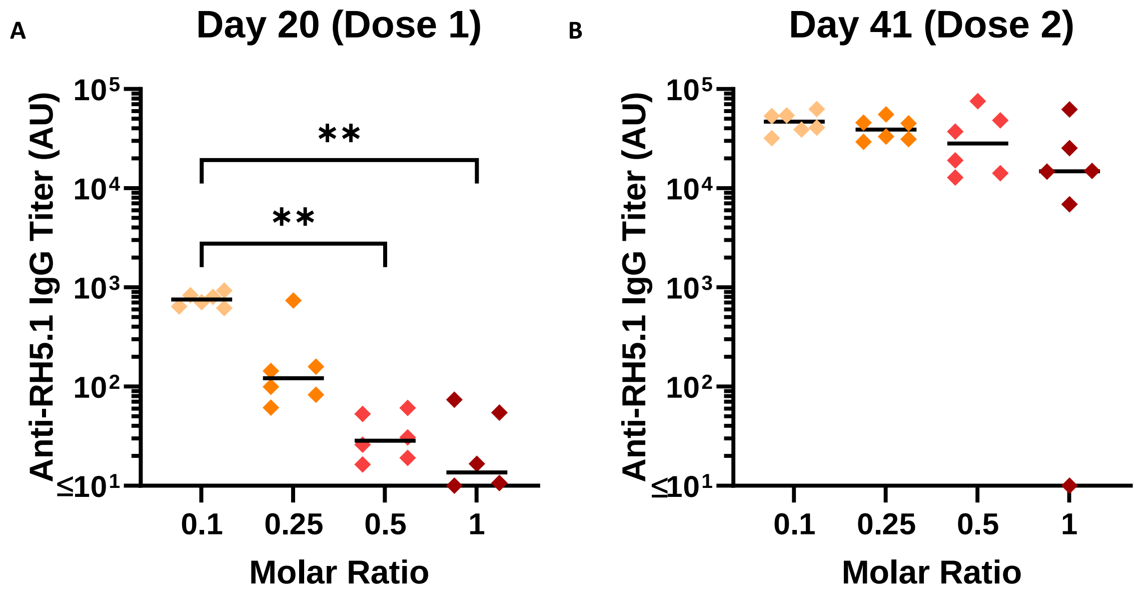


**Figure S4. Immunogenicity testing of the RH5.2-VLP vaccine produced with different conjugation efficiencies.**

BALB/c mice (N=6 per group) were immunized intramuscularly with three doses of RH5.2-VLP, produced using the indicated molar ratios of RH5.2-ST to HBsAg-SC (0.1:1, 0.25:1, 0.5:1 and 1:1), on days 0, 21 and 42. Dosing was adjusted in each case to deliver the same molar amount of RH5.2 antigen (10 ng); total RH5.2-VLP dose = 232, 52, 40 and 23 ng, respectively. All vaccines were formulated in Matrix-M™ adjuvant. Anti-RH5 (full-length RH5.1) IgG titers were measured in the serum by ELISA after (**A**) dose 1 at day 20, and (**B**) dose 2 at day 41. Each point represents a single mouse and the line the median. Analysis using Kruskal-Wallis test with Dunn's multiple comparison test across the four groups; ***P* < 0.01.
